# The LDB1 complex co-opts CTCF for erythroid lineage specific long-range enhancer interactions

**DOI:** 10.1101/128082

**Authors:** Jongjoo Lee, Ivan Krivega, Ryan K. Dale, Ann Dean

**Author notes:** Corresponding author: Building 50, Room 3154 50 South Drive, MSC 8028 Bethesda, MD 20892.

## Abstract

Lineage-specific transcription factors are critical for long-range enhancer interactions but direct or indirect contributions of architectural proteins such as CTCF to enhancer function remain less clear. The LDB1 complex mediates enhancer-gene interactions at the β-globin locus through LDB1 self-interaction. We find that a novel LDB1-bound enhancer upstream of *carbonic anhydrase 2* (*Car2*) activates its expression by interacting directly with CTCF at the gene promoter. Both LDB1 and CTCF are required for enhancer-*Car2* looping and the domain of LDB1 contacted by CTCF is necessary to rescue *Car2* transcription in LDB1 deficient cells. Genome wide studies and CRISPR/Cas9 genome editing indicate that LDB1-CTCF enhancer looping underlies activation of a substantial fraction of erythroid genes. Our results provide a mechanism by which long-range interactions of architectural protein CTCF can be tailored to achieve a tissue-restricted pattern of chromatin loops and gene expression.

## INTRODUCTION

Enhancers are regulatory elements that act over long distances to activate transcription of target genes, thereby defining the unique transcriptomes of cells and tissues. The physical interaction between enhancers and their target genes is crucial for this activity and the cell type specificity of the contacts depends on lineage-restricted activators (de Laat and Duboule, 2013; Plank and Dean, 2014; Gorkin et al., 2014). Enhancer-gene interactions occur primarily within topologically associating domains (TADs) that spatially constrain them (Dixon et al., 2012; Shen et al., 2012; Nora et al., 2012). The architectural protein CCCTC-binding factor (CTCF) and its frequent partner cohesin contribute to TAD organization at multiple levels. CTCF sites are enriched at TAD borders, however, the vast majority of CTCF sites occur within TADs (Phillips-Cremins et al., 2013; Merkenschlager and Odom, 2013; Ong and Corces, 2014; Gorkin et al., 2014). Moreover, genome wide studies suggest a role for architectural proteins, including CTCF and cohesin, and Mediator in spatially connecting enhancers and the genes they regulate within TADs (Handoko et al., 2011; Phillips-Cremins et al., 2013; Heidari et al., 2014; Rao et al., 2014; Ing-Simmons et al., 2015; Bouwman and de Laat, 2015). Whether CTCF plays a direct or indirect role in enhancer-gene interactions and how such a role can be reconciled with the strong tissue-specificity of enhancers are critical unanswered questions.

Focused studies have provided some clues as to how CTCF might participate in locus specific long range interactions. For example, the interaction of CTCF with pluripotency factor OCT4 in ES cells is central to long range interactions involved in X chromosome inactivation (Donohoe et al., 2009). The role of CTCF in enhancer-gene looping has also been studied at select gene loci.

In the *Ifng* locus, CTCF promotes loops between its sites within and near *Ifng* with distant enhancer sites occupied by the lineage-specific transcription factor T-BET (Sekimata et al., 2009). However, CTCF and T-BET proteins were not observed to interact. In the *Myb* and *Tal1* loci, interspersed CTCF sites and enhancers occupied by LDB1 form complex looped conformations when the genes are actively transcribed in erythroid cells (Stadhouders et al., 2012; Zhou et al., 2013). The mechanisms underlying these tissue specific enhancer looping interactions remain unclear.

LDB1 is a transcription co-factor that is essential for long-range interaction of the β-globin locus control region (LCR) enhancer with β-globin genes, which is required for their activation (Song et al., 2007; Krivega et al., 2014). LDB1 does not bind DNA directly but is recruited to compound E box/GATA elements in the LCR and β-globin promoter via a multi-component complex that includes erythroid DNA-binding factors GATA1 and TAL1, and bridging protein LMO2. LDB1 interacts with LMO2 through the C-terminal *lim interaction domain*, while interaction between LDB1 N-terminal self-dimerization domains supports long range LCR/gene interaction (Deng et al., 2012; Deng et al., 2014; Krivega et al., 2014).

Genome-wide studies suggest that LDB1 complexes function broadly at enhancers to activate erythroid genes (Fujiwara et al., 2009; Yu et al., 2009; Kassouf et al., 2010; Soler et al., 2010; Li et al., 2013; Mylona et al., 2013). However, microarray studies in MEL cells with reduced LDB1, and ChIP-seq studies had revealed numerous genes that are positively regulated by LDB1 but whose promoters, unlike β-globin, are not occupied by the LDB1 complex. Among these is *carbonic anhydrase 2* (*Car2*), one of the most strongly down-regulated genes upon reduction of LDB1 in erythroid cells (Song et al., 2012). The genes encoding *carbonic anhydrases 2* and *3* are clustered together in mouse and human on chromosomes 3 and 8, respectively, with *Car1* located about 100 kb distant. These soluble anhydrases are members of a large family of proteins that function to exchange CO_2_ and O_2_ (Edwards et al., 2000). *Car3* is expressed predominantly in smooth muscle cells, while *Car1* and *Car2* are expressed predominantly in erythroid cells. *Car2* is highly transcribed in fetal liver erythroid cells (ENCODE Project Consortium, 2014) and is activated during erythroid maturation in MEL cell and G1E cell model systems, similar to numerous genes that comprise the mature erythroid transcriptome (Welch et al., 2004; Song et al., 2012).

Here we show that LDB1-CTCF-mediated enhancer looping underlies activation of numerous erythroid genes. Using the *Car2* gene as an example, we find that LDB1 bound to an upstream enhancer and CTCF bound to the gene promoter interact physically and functionally to mediate activation of *Car2* in erythroid cells. Moreover, we identify a subset of CTCF-occupied genes that loop to LDB1-bound known or putative erythroid enhancers. CRISPR/Cas9 deletion of select candidate enhancers compromises gene transcription, validating enhancer function and generalizing the importance of LDB1-CTCF interaction in enhancer looping. Our results reveal direct participation of CTCF in tissue-specific long-range enhancer interactions and in establishment of the erythroid transcriptome.

## RESULTS

### LDB1 sites upstream of *Car2* function as an enhancer of *Car2* expression

During erythroid maturation, *Car2* is activated (Welch et al., 2004; Song et al., 2012) in parallel with β-globin, although to very much lower levels, while key regulators such as GATA1 and LMO2 maintain similar levels of transcription (Figure 1A). ChIP and deep sequencing revealed that LDB1 and complex members GATA1 and TAL1 occupy a pair of intergenic sites located -8 and -9 Kb upstream of the *Car2* promoter and downstream of *Car3* (inactive in erythroid cells) in mouse bone marrow cells (Li et al., 2013), primary mouse erythroid cells (Yu et al., 2009) and MEL cells (Soler et al., 2010; Song et al., 2012)(Figure 1B). Moreover, these sites display prominent enhancer marks such as H3K27ac, H3K4me1 and p300, making them strong candidate regulatory elements.

**Figure 1.**
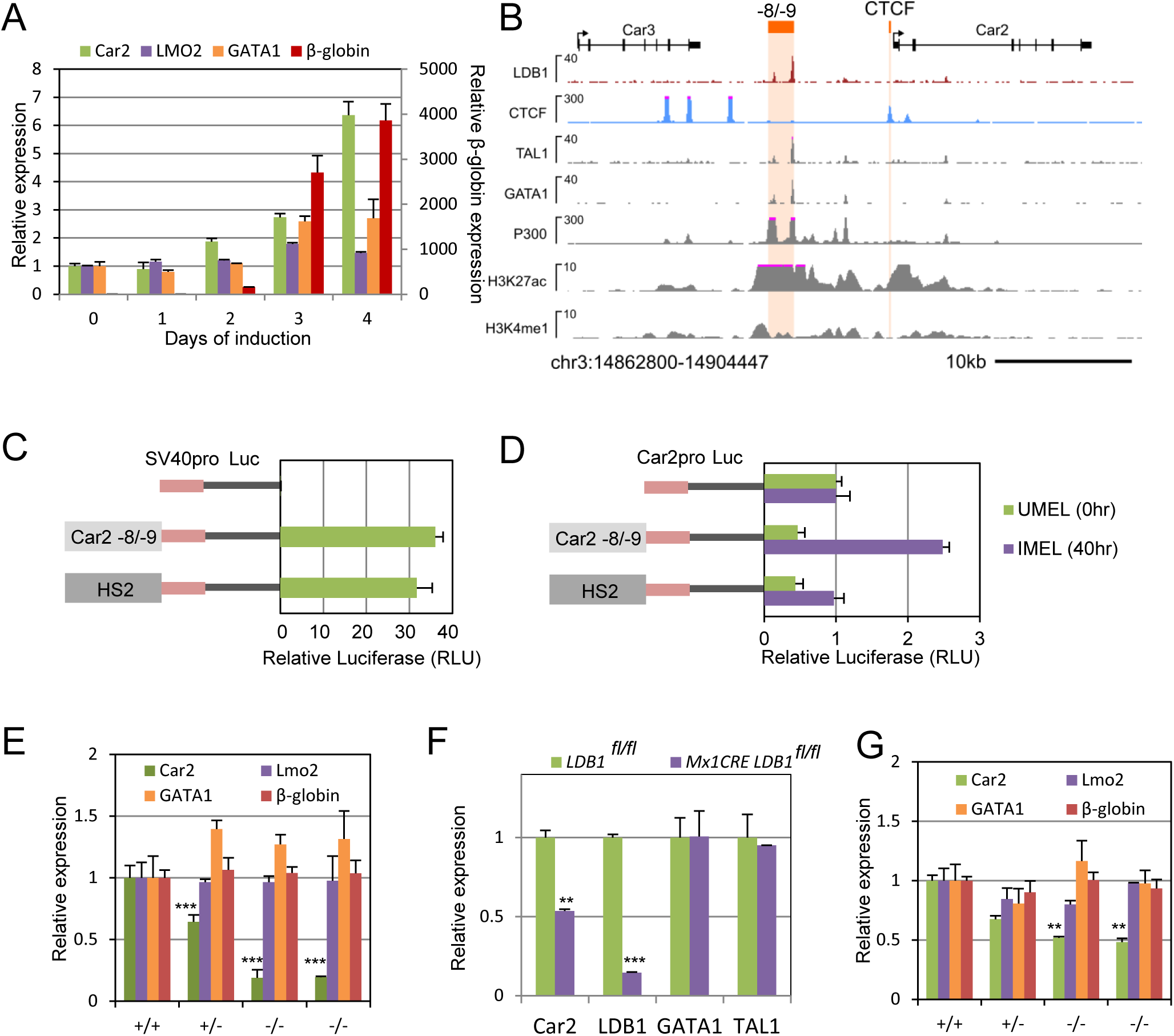
An upstream, LDB1 enhancer controls Car2 expression. (A) *Car2, Lmo2, Gata1* (left axis) and *β-globin* (right axis) expression during differentiation of MEL cells with DMSO. Values on day 1 were set to 1. (B) Genome browser view of the *Car2* and *Car3* genes with protein occupancy and histone modification data for uninduced cells. ChIP-seq tracks are from published data (Soler et al., 2010; ENCODE Project Consortium, 2014). LDB1 sites and *Car2* promoter CTCF site are highlighted in orange. (C, D) Transient reporter assays in K562 cells for the constructs shown in C or in MEL cell for the constructs shown in D. Relative luciferase activity is plotted in (D). Value for the enhancerless construct set to 1. (E) Expression of *Car2* and control genes in enhancer-deleted clones. Two homozygous deletion clones are shown. WT value was set to 1. (F) Expression of the *Car2, Ldb1, Gata1* and *Tal1* genes in IFN-β-treated E14.5 *Ldb1*^fl/fl^ fetal liver cells with and without Mx1Cre. Expression level in E14.5 *Ldb1*^fl/fl^ without Mx1CRE cells was set to 1. (G) Expression of *Car2* and control genes in *Car2* CTCF site-deleted clones. Two homozygous deletion clones are shown.WT value was set to 1. See also Figures S1 and S2.

To ask if these LDB1 sites function to enhance transcription of *Car2*, we first carried out luciferase reporter assays in which the *Car2* -8/-9 sequences or the well-known LCR HS2 enhancer of the β-globin locus were cloned upstream of the SV40 promoter (Figure 1C). In K562 cells, a human erythroid cell line, the *Car2* locus LDB1 sites showed equivalent enhancer activity to LCR HS2 compared to a vector with only a promoter. In MEL cells, the activity of the -8/-9 kb region was tested using a 1 kb region encompassing the transcription start site of *Car2* as promoter (Figure 1D). Upon induction with DMSO the -8/-9 region increased luciferase activity about 2-3 fold, which was greater than the effect seen with the HS2 enhancer.

Next, we deleted sequences encompassing the *Car2* -8/-9 LDB1 sites by using CRISPR/Cas9 mediated genome editing in MEL cells (Cong et al., 2013) (Table S1, Figure S1). Mono- or bi-allelic deletion reduced *Car2* transcription in induced MEL cells in a dose-dependent fashion (Figure 1E). An additional single gRNA more specifically targeted LDB1 complex binding at -9 kb by deleting the GATA1 site. Transcription of *Car2* was reduced by half, an outcome suggesting that -8 and -9 kb LDB1-bound complexes contribute equally to activation of Car2 (Figure S2A). Transcription of other genes required for erythroid differentiation such as *Gata1* and *Lmo2* were not significantly affected, nor was differentiation altered as judged by normal induction of β-globin mRNA (Figure 1E). Together, these data provide compelling evidence for an upstream LDB1 *Car2* enhancer active in erythroid cells that is LDB1 dependent.

To investigate a role for LDB1 in *Car2* expression *in vivo*, we took advantage of a mouse model of conditional *Ldb1* deletion (Li et al., 2010; Krivega et al., 2014). *Cre* expression in E14.5 fetal livers, consisting primarily of erythroid cells, results in >50% excision of *Ldb1* (Krivega et al., 2014). Under these conditions, both *Ldb1* and *Car2* expression are significantly reduced (Figure 1F). These *in vivo* data support the idea that LDB1 is essential for *Car2* activation during erythroid differentiation.

In contrast to β-globin, the Car2 promoter is occupied by CTCF but not by the LDB1 complex (Figure 1B). To address the function of the CTCF binding in the *Car2* promoter, we used CRISPR/Cas9 to delete promoter sequences encompassing two adjacent high-scoring matches to the CTCF motif (JASPAR database (Mathelier et al., 2015) (Table S1, Figure S1). The deleted region was unoccupied by factors besides CTCF for which ChIP-seq data are available from ENCODE for erythroblasts and MEL cells. *Car2* transcription was down-regulated in induced MEL cells in a dose-dependent fashion by mono- or biallelic deletion of the promoter CTCF occupancy region (Figure 1G), while control genes were normally transcribed, indicating normal erythroid differentiation. An additional single gRNA disrupting only the two CTCF motifs and the 21 bp between them similarly reduced Car2 transcription strongly (Figure S2B). These data show that the promoter CTCF interaction is an important component of *Car2* transcription activation.

### *Car2* is regulated by LDB1 and CTCF through interaction between the *Car2* promoter and upstream enhancer

LDB1-mediated enhancers are predicted to activate target genes by long-range interactions (Song et al., 2007; Krivega et al., 2014). To examine chromosome folding in the vicinity of *Car2*, we carried out chromatin conformational capture (3C) across 40 kb of mouse chromosome 3 using the *Car2* -8/-9 kb enhancer as the viewpoint (Figure 2A). Reverse primers were located in each fragment generated by the frequent cutter BstY1, allowing a view of enhancer-promoter interactions in the context of all enhancer contacts formed. Compared to uninduced cells, long-range interactions of the *Car2* enhancer were evident in induced cells, with peaks corresponding to CTCF sites in the *Car2* promoter and first intron. The enhancer also contacted two upstream CTCF sites, one of which is within the body of the neighboring *Car3* gene (silent in erythroid cells). No other contacts were observed for the enhancer over 150 kb surrounding the Car2 locus (Figure S3). Biallelic deletion of the CTCF site in the *Car2* promoter (Figure 1G) abolished long-range interactions seen in induced cells between the *Car2* enhancer and gene (Figure 2A).

**Figure 2.**
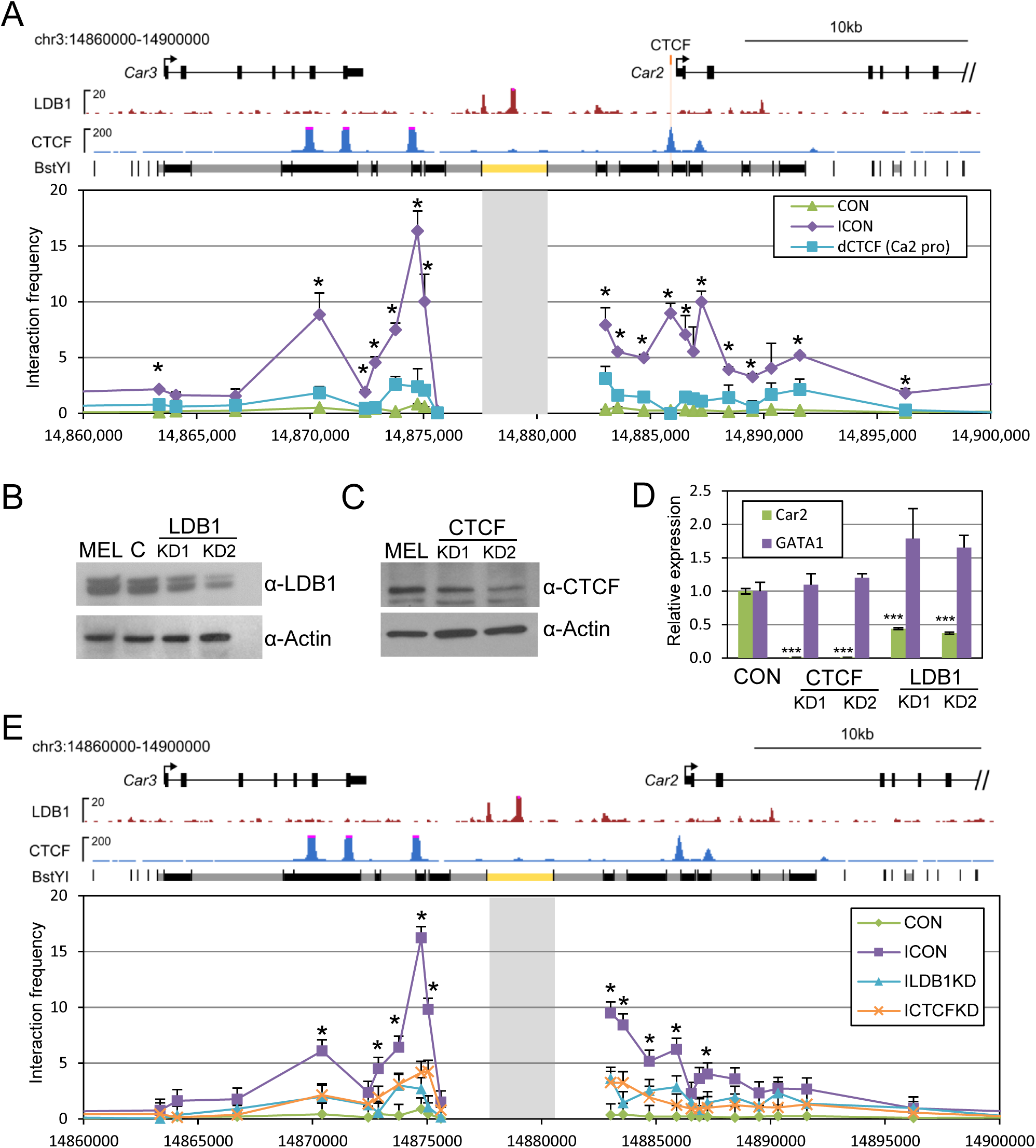
LDB1 and CTCF are required for chromatin looping between the *Car2* enhancer and gene. (A) Interaction frequency determined by 3C between locations across *Car2*—*Car3* using the *Car2* enhancer as the anchor (shaded). BstY1 restriction sites and interrogated fragments (alternately shown in black or gray) are shown across the top. The anchor fragment is indicated in yellow. CON, control cells; ICON, induced control cells; dCTCF, induced representative clone with the *Car2* promoter CTCF site deleted. *, P<0.05 for comparison between induced control MEL cells and induced dCTCF cells. (B, C) Western blots illustrate protein reduction in representative stable MEL cell clones after shRNA against LDB1 (B) or CTCF (C). C, control scrambled shRNA vector. KD, knockdown. Actin served as control. (D) Relative expression of *Car2* and *Gata1* in representative KD clones. CON, control scrambled shRNA vector. (E) Interaction frequency determined by 3C between locations across *Car2*—*Car3* using the *Car2* enhancer as the anchor (shaded). Details are the same as for panel A. CON, control, ICON; induced control. KD, knockdown. *, P<0.05 for comparison between induced control MEL cells and both induced LDB1KD MEL cells and induced CTCFKD MEL cells. See legend to Figure 1B for sources of tracks shown. See also Figure S3.

We next tested whether the transcription of *Car2* is regulated by LDB1 and CTCF by using shRNA mediated reduction of these proteins. In stable, LDB1 knockdown (KD) MEL cell clones, numerous genes required for erythroid maturation are transcribed normally, although β-globin is not activated when the cells are induced (Song et al., 2007; Song et al., 2010; Li et al., 2010; Krivega et al., 2014). Clones with 2-3-fold reduced expression of LDB1 or CTCF had substantially lower levels of these proteins than WT cells (Figure 2B and C). *Car2* expression was strongly reduced in the KD clones while, as a control, *Gata1* expression was not significantly affected (Figure 2D). LDB1 KD did not affect CTCF transcription or protein levels and vice versa (Figure S4). When either LDB1 or CTCF was reduced, chromatin looping in induced cells between the -8/-9 LDB1 enhancer sites and *Car2* was compromised (Figure 2E). These results indicate that *Car2* activation is associated with chromatin loop formation between an LDB1-occupied *Car2* enhancer and the CTCF-occupied *Car2* promoter and that both LDB1 and CTCF are important to *Car2* enhancer long-range interactions and to *Car2* activation.

### LDB1 directly interacts with CTCF

Inspection of mouse ENCODE data indicates that CTCF and its frequent partner cohesin jointly occupy the *Car2* promoter CTCF site. We also observed that KD of the SMC3 cohesin component strongly reduced *Car2* transcription (not shown). Thus, we considered whether LDB1 and CTCF/cohesin might interact to activate *Car2* transcription. We included the Mediator co-activator complex in the analysis as it is known that cohesin interacts with mediator to loop enhancer and promoter regions together for gene activation in ES cells (Kagey et al., 2010).

We performed co-immunoprecipitation experiments with nuclear extracts of induced and uninduced MEL cells. As expected, based on cohesin interactions with Mediator and with CTCF (Kagey et al., 2010; Xiao et al., 2011), MED1 and cohesin loading factor NIPBL can immunoprecipitate CTCF and cohesin component SMC1 (Figure 3A). Similarly, antibodies to GATA1 immunoprecipitate CTCF and MED1 (Stumpf et al., 2006; Manavathi et al., 2012) and SMC1, possibly indirectly (Xiao et al., 2011). Consistent with GATA1 principally functioning as part of the LDB1 complex (Li et al., 2013), LDB1 antibodies also immunoprecipitate CTCF and SMC1, although not MED1.

**Figure 3.**
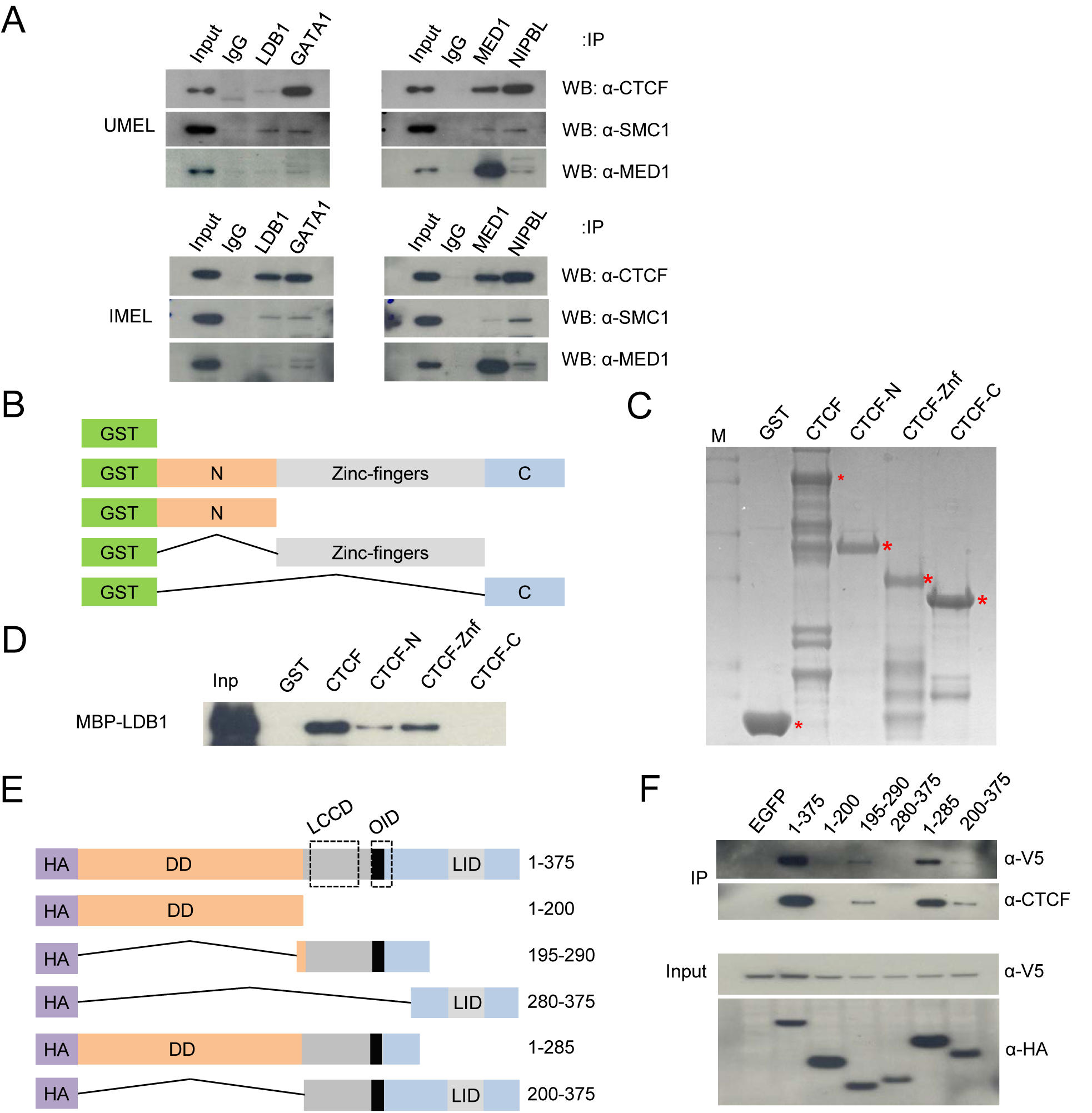
LDB1 and CTCF interact directly. (A) Co-IP of endogenous proteins was performed using MEL cell nuclear extracts treated with (IMEL) or without (UMEL) DMSO for 4 days. Antibodies to LDB1, GATA-1, MED1 or NIPBL were used for immunoprecipitation and blots developed with antibodies to CTCF, SMC1 and MED1. The input lane contains 5% of the immunoprecipitated material. (B) Diagram of GST-tagged versions of CTCF tested. (C) Coomassie blue stained gel illustrating production of the CTCF proteins shown in B in bacteria (labeled with an asterisk). (D) Western blot analysis of interaction *in vitro* between full length and truncated forms of CTCF and MBP-tagged LDB1developed with antibodies to the MBP tag. (E) Diagram of HA-tagged versions of LDB1 tested for interaction with CTCF. LCCD, OID and LIM domains of LDB1 are indicated (see text). (F) Western blot developed with antibodies to either the V5 tag or to CTCF of the input and material immunoprecipitated (IP) by full length or truncated forms of HA-tagged LDB1. The successful production of each of the proteins in 293T cells is shown below in a blot developed with antibodies to the HA tag. See also Figure S4.

Interestingly, the LDB1-CTCF immunoprecipitation is highly enriched in induced MEL cells. Levels of CTCF and LDB1 are unchanged upon induction of MEL cells (Song et al., 2007) (Figure S4). The difference may reflect post-translational modification of CTCF and/or association of additional proteins that may favor interaction in induced cells or antagonize interaction in uninduced cells. Overall, the results suggest that CTCF may be important for enhancer looping in erythroid cells and raise the possibility that the LDB1 complex specifyies the enhancers to be involved.

To further investigate potential interaction of CTCF and LDB1, we carried out *in vitro* GST pull-down assays using *E. coli* expressed full-length GST-tagged CTCF and deletion constructs including N-terminal, Zinc-finger or C-terminal domains (Figure 3B, C). Figure 3D shows that LDB1 interacted with full length CTCF and with the CTCF Zinc finger domain, a common interaction module for CTCF. LDB1 did not interact with the CTCF C-terminal region LDB1 and only weakly with the N-terminal domain.

Using a similar approach, N-terminal HA tagged deletion mutants of LDB1 (Figure 3E) were expressed along with V5-tagged CTCF in 293T cells. The minimal region of LDB1 required for CTCF interaction contained the *LDB1/Chip conserved domain* (LCCD) and NLS (van Meyel et al., 2003), and the *other interacting domain* (OID) through which *Drosophila* Chip interacts with the insulator protein Su(Hw) (Torigoi et al., 2000) (Figure 3F). Because GATA1 is not expressed in 293T cells, we conclude that it is not necessary for interaction between CTCF and LDB1. These results support the idea that LDB1 and CTCF interact through a specific domain in each protein

### The LDB1 CTCF-interacting module is necessary and sufficient for Car2 activation

To test the importance of the LDB1 LCCD domain for interaction with CTCF, we carried out loss and gain of function studies. First, we used CRISPR-Cas9 to target LDB1 by deletion of exon 9 in MEL cells (Figure 4A, B). Stable clones were obtained in which LDB1 was undetectable by western blot analysis and *Car2* protein was, as expected, greatly reduced (Figure 4C). *Car2* expression was undetectable in LDB1 KO cells ectopically expressing a control EGFP vector but was rescued upon expression of HA-tagged LDB1 (Figure 4D). However, HA-tagged LDB1 missing the LCCD domain failed to rescue Car2 expression over levels seen with a mock transfection, supporting the necessity of the LCCD for long-range function of the *Car2* enhancer.

**Figure 4.**
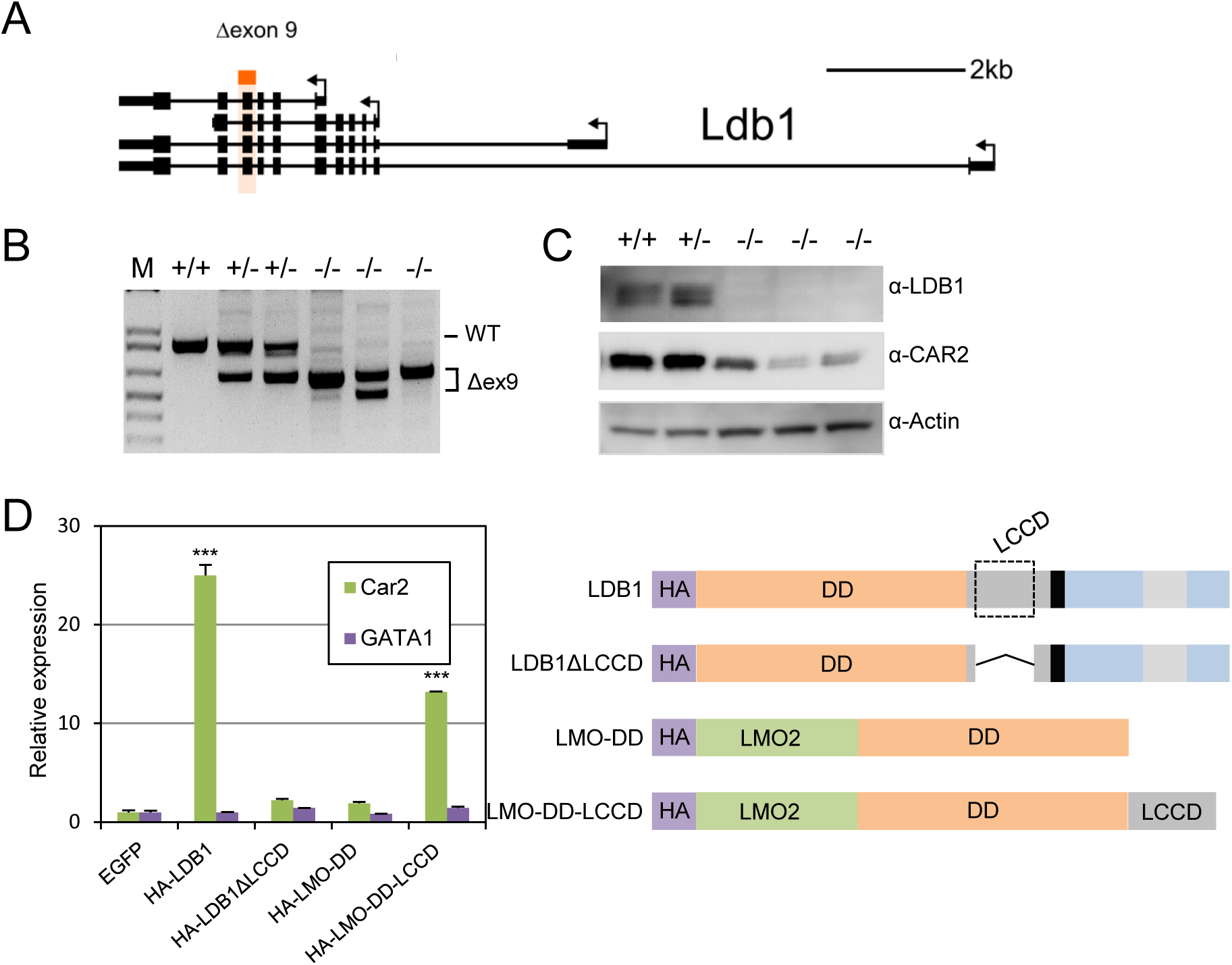
LDB1 LCCD is necessary and sufficient to interact with CTCF. (A) Gene model for *Ldb1* with exon 9 boxed in red. (B) PCR validation of *Ldb1* exon 9 deletion using flanking primers. Representative WT and mono- or biallelically deleted clones are shown. (C) Western blot of cell extracts from representative WT and mono- or biallelically deleted clones are shown. Blots were developed with antibodies to LDB1 or CAR2 and actin served as control. (D) RT-qPCR with RNA extracted from LDB1 KO MEL cells ectopically expressing full length HA-LDB1, HA-LDB1ΔLCCD, HA-LMO-DD or HA-LMO-DD-LCCD or EGFP as control. Error bars indicate SEM; n=3 biological replicates.

LDB1 KD cells also fail to express β-globin but transcription can be rescued by ectopic expression of full length LDB1 (Krivega et al., 2014). β-globin transcription can also be rescued in KD cells by expression of a fusion of the LDB1 dimerization domain with LMO2 (LMO-DD), indicating the necessity and sufficiency of the DD for β-globin rescue (Krivega et al., 2014). LMO-DD fails to rescue *Car2* expression in LDB1 KO cells Figure 4D). However, inclusion of the LCCD in the fusion protein (LMO-DD-LCCD) resulted in significant rescue of *Car2* expression. These loss and gain of function experiments establish the role of the LDB1 LCCD in long-range activation of *Car2*. Since all rescue constructs contained the LDB1 DD, we cannot rule out a direct or indirect contribution of the DD to *Car2* rescue. Overall, we conclude that LDB1-CTCF interaction provides an unexpected mechanism to recruit CTCF into an erythroid lineage specific enhancer looping function.

### LDB1-bound enhancers loop to genes that are occupied by CTCF

We next considered whether the example of *Car2* long range regulation by the LDB1 complex and CTCF might be of more general importance to enhancer looping and function in erythroid cells. Notably, the fetal γ-globin repressor *Bcl11a* (Xu et al., 2010) loops to and is activated by intronic LDB1-occupied enhancers but the *Bcl11a* promoter is occupied by CTCF rather than LDB1, similar to *Car2* (Bauer et al., 2013; ENCODE Project Consortium, 2014). Likewise, the *Myb* gene has no sites of LDB1 occupancy but several LDB1 occupied putative enhancers appear to loop to the promoter/ first intron that contains a CTCF site (Stadhouders et al., 2012).

To gain further insight into the genome wide erythroid enhancer repertoire we used ENCODE ChIP-seq data from uninduced MEL cells and the ChromHMM algorithm (Ernst et al., 2011) to build a set of hidden Markov models (https://doi.org/10.5281/zenodo.439534 and see Supplemental Material). We identified a 6-state model that used H3K4me1 and H3K27ac (enhancer marks), H3K4me3 and H3K36me3 (active gene marks), H3K27me3 (repressed chromatin) and DNase-seq (regulatory regions) as the model with the most readily interpretable biological states (Figure S5A, and see supplemental methods). The model called 48,041 enhancers in erythroid cells of which 7,765 (16%) were occupied by LDB1 (Figure S5B). This is likely to represent a sub-set of LDB1 complex-occupied enhancers, since when we used GATA1 Chip-seq data as a proxy for LDB1 complex occupancy, 53% of enhancers were scored as positive (not shown).

To obtain stringent genome-wide identification of genes connected to these predicted enhancers, we intersected these data with promoter capture Hi-C results from erythroid cells, a powerful means of determining relevant functional enhancer-gene pairs (Schoenfelder et al., 2015). We found that 84% of our called enhancers looped to at least one gene and 88% of the enhancers occupied by LDB1 were so engaged (Figure S5B), providing functional validation of our called enhancers. Thus, overall, enhancers in erythroid cells are strongly linked to genes by long-range interactions.

Figure 5A illustrates our ChromHMM model prediction of the known +23 kb *Runx1* enhancer (Nottingham et al., 2007). The LDB1 complex occupies the enhancer and gene promoter, similar to the configuration in the β-globin locus (Song et al., 2007), and long range interaction occurs between them (Schoenfelder et al., 2015). Long range interactions were also observed between LDB1-occupied putative enhancers and genes not occupied by LDB1 (Figure 5B, C). For example, the *Cpeb4 Plcl2* genes interact with called distant enhancer sites occupied by LDB1. *Cpeb4* and *Plcl2* are not themselves occupied by LDB1 but have CTCF sites either in the promoter (*Plcl2*, similar to *Car2*) or in the first intron (*Cpeb4*, similarly to *Myb*).

**Figure 5.**
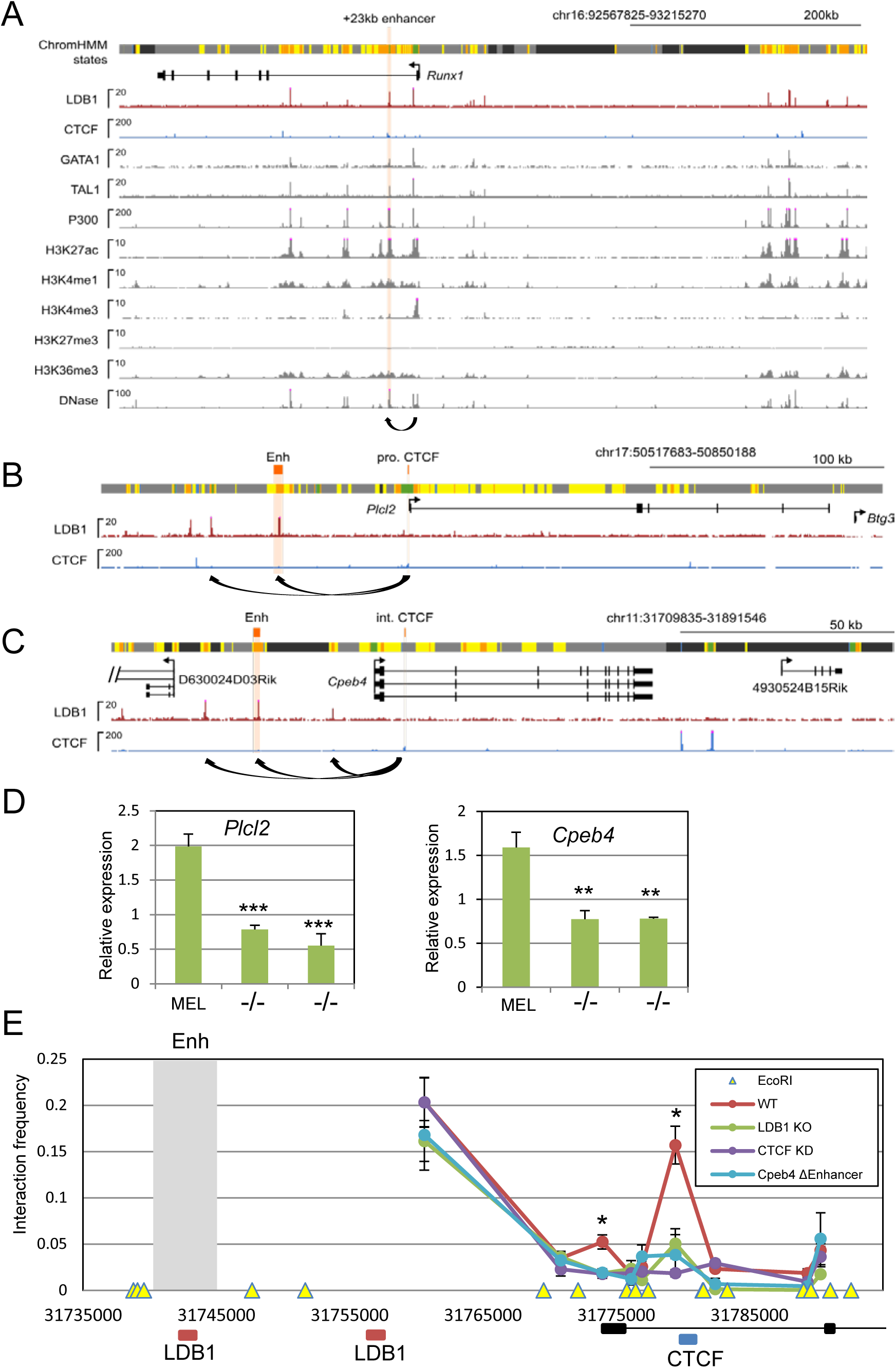
LDB1 and CTCF regulation of erythroid genes by enhancer-promoter looping. (A) Genome browser view of the *Runx1* locus. Shown are positions of ChromHMM states color-coded as in Figure S5A. Gene model is from RefSeq, and signals are from data used in the models, plus additional tracks for GATA1, TAL1, and P300. Enhancer and CTCF regions discussed in the text are highlighted in orange (pro, promoter; int, intron; enh, enhancer). (B) Genome browser view of *Plcl2* and (C) *Cpeb4*. For each view, LDB1 and CTCF occupancy are shown. Capture-C looping interactions reported by Schoenfelder *et al* (Schoenfelder et al., 2015) are shown as curved black arrows. Enh, putative enhancer; int, intron; pro, promoter. (D) Graphs show gene expression for representative clones after CRISPR/Cas9 mono- or biallelic deletion of the indicated putative enhancer compared to WT cells. Two homozygous deletion clones are shown. (E) Interaction frequency determined by 3C between locations across the *Cpeb4* locus using a predicted enhancer as the anchor (shaded). Eco RI restriction sites are indicated by the yellow triangles across the bottom. *Cpeb4*ΔEnhancer, representative clone with the *Cpeb4* enhancer deleted. LDB1 and CTCF tracks in panels A-C are from published sources (Soler et al., 2010; ENCODE Project Consortium, 2014). In panels D and E, error bars indicate SEM; n=3 biological replicates. * = p<0.05, ** = p<0.01 and *** = p<0.001 by Student’s T test compared to uninduced WT MEL cells. See also Figure S5.

To ask whether the LDB1-occupied sites in contact with these genes are *bona fide* enhancers, we used the CRISPR/Cas9 genome editing approach (Table S1). Bi-allelic deletion of LDB1- occupied putative intergenic enhancers that looped to *Cpeb4* or *Plcl2* reduced expression 3-4-fold but did not eliminate it entirely (Figure 5D). In both these loci, additional LDB1-occupied potential enhancers engage in looping interactions with the genes, likely accounting for additional regulatory influences, consistent with observations genome wide of multiple enhancers interacting with individual genes (Sanyal et al., 2012; Jin et al., 2013; Schoenfelder et al., 2015).

Chromosome conformation capture (3C) validated the capture Hi-C identification of loops between *Cpeb4* and putative LDB1-occupied upstream enhancers (Figure 5C, E). Deletion of one of these enhancers resulted in significant loss of interaction with the CTCF site in intron 1, which is consistent with the transcription reduction (Figure 5E). In addition, shRNA-mediated reduction of CTCF or KO of LDB1 also reduced *Cepb4* long range interactions with the enhancer. These deletion and KO studies support the idea that LDB1-CTCF interaction underlies looping and enhancer activity at select loci in erythroid cells.

### Erythroid Genes are preferentially engaged by LDB1 bound enhancers in erythroid cells

Within the set of genes queried for long range interactions (Schoenfelder et al., 2015), we compared a literature-curated set of 775 erythroid genes (http://doi.org/10.5281/zenodo.189503 and see Supplemental Material) to the remaining set of annotated genes (24,806, here after “other” genes) and classified genes into subsets depending on enhancer contact. Erythroid genes were enriched for interactions with least one enhancer (p=4.1e-66; odds ratio 6.8, Fisher’s exact test) and for interactions with an LDB1-bound enhancer (p=1.6e-78; odds ratio 4.3, Fisher’s exact test) compared to other genes (Figure S5C).

Almost all (94%) of the erythroid genes looped to an enhancer and of these, 582 (80%) looped to an LDB1-bound enhancer, supporting the idea that LDB1-bound enhancers are the predominant activators of erythroid genes (Fujiwara et al., 2009; Yu et al., 2009; Kassouf et al., 2010; Soler et al., 2010; Li et al., 2013; Mylona et al., 2013) (Figure S5C). However, of the LDB1-bound enhancers that looped to a gene, only 24% were looped to an erythroid gene (Figure S5B) and the remainder contacted other genes, suggesting that LDB1 enhancers have more broad functions in erythroid cells than had been previously appreciated. We note that the set of erythroid genes is not exhaustive and there are likely to be erythroid genes within the set of ‘other’ genes.

Alternatively, ‘other’ genes may share certain components of the enhancer mechanisms we propose here for erythroid genes.

Erythroid genes were expressed at significantly higher levels than other genes (Figure S5C) (9.3 vs 1.8 average TPM, p=5.4e-47, Mann-Whitney U test). Erythroid genes also looped to multiple LDB1-bound enhancers significantly more frequently than other genes (2.6 vs 1.0 mean enhancers per gene, p=7.0e-88, Mann-Whitney U test) (Figure S5D). For example, the highly-connected erythroid gene, *Epor*, interacts with seven LDB1-bound enhancers, some over 150 kb away (Figure S5E). The number of enhancers per erythroid gene increased further when GATA1 was used as a proxy for LDB1 complex occupancy (not shown). Taken together, these results indicate that in erythroid cells, erythroid genes are preferentially contacted by enhancers compared to other genes and that among these contacts, LDB1-occupied enhancers are strongly enriched.

### Long range communication to erythroid genes by LDB1 occupied enhancers

Interestingly, both LDB1 and CTCF are bound at the multiply looped *EpoR* promoter (Figure S5E). This result raises the question of the contributions, both individually and together, of LDB1-CTCF interactions described here and the previously reported dimerization of LDB1 (Krivega et al., 2014) within the erythroid enhancer contact landscape.

Figure 6A (and see Figure S5C) compares LDB1 and CTCF occupancy at erythroid genes and ‘other’ genes that were looped to at least one LDB1-bound enhancer in erythroid cells. Of the 582 erythroid genes that contacted an LDB1-bound enhancer, 84 (14%) had LDB1 but not CTCF at their promoter (similar to β-globin), while 210 (36%) had CTCF but not LDB1 (similar to *Car2)*, at their promoter and 178 (31%) were occupied by both proteins (similar to *EpoR)*. Promoters with LDB1 alone or with LDB1 and CTCF are strongly enriched among erythroid genes compared to other genes (odds ratio=3.2, p=1.3e-16, Fisher’s exact test; odds ratio=3.8, p=2.9e-37, Fisher’s exact test, respectively). In contrast, CTCF without LDB1 appears commonly at the promoters of both erythroid genes and other genes (odds ratio=0.91, p=0.31, Fisher’s exact test).

**Figure 6.**
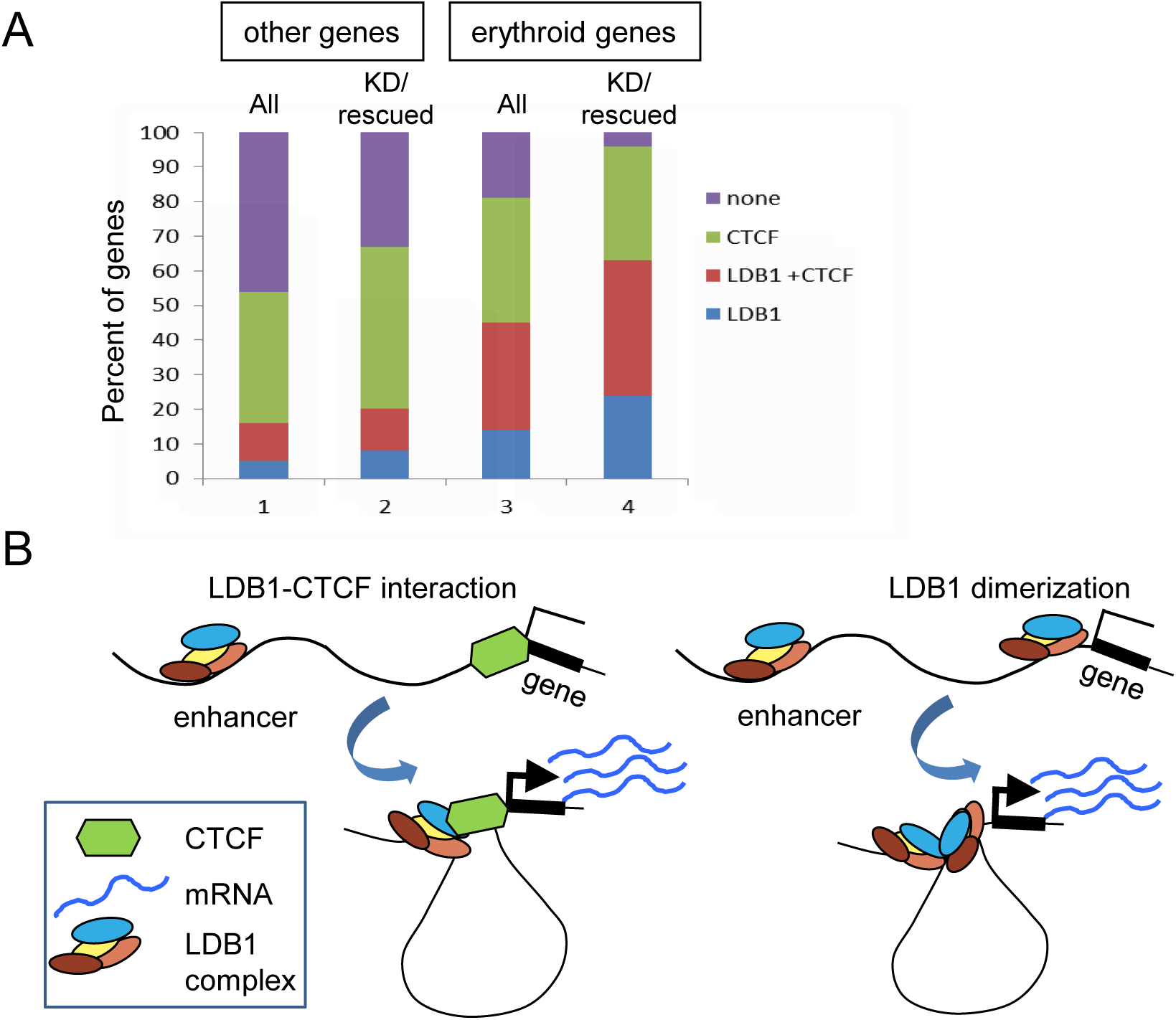
Erythroid gene occupancy by LDB1 and CTCF is common. (A) Genes are considered in two groups: erythroid genes and other genes. For each group the percent of genes whose promoters are occupied by LDB1, CTCF, both proteins or neither protein is plotted. Each group is considered as a whole (all) and the group whose expression is reduced upon LDB1 KDknockdown and rescued by expression of full length LDB1 in MEL cells is considered separately (KD/rescued). (B) Models depicting long range interaction between an LDB1-bound enhancer and a CTCF or LDB1 occupied gene to activate expression.

Comparison with RNA-seq analyses revealed that 46 erythroid genes looped to an LDB1- occupied enhancer were reduced upon LDB1 KD and rescued by expression of full length LDB1(Krivega et al., 2014). There were 198 ‘other’ genes in this repressed and rescued group, which is understandably a larger number since most LDB1-bound enhancers loop to non-erythroid genes (Figure S5C). The promoters of the LDB1-enhancer-looped erythroid genes were occupied by LDB1 alone (11 genes, 24%), CTCF alone (15 genes, 33%) or both (18 genes, 39%). Only 2 (4%) erythroid gene promoters had neither protein, in contrast to 65 ‘other’ genes (33%). This suggests that ‘other’ genes use different proteins to contact LDB1-occupied enhancers than do erythroid genes. We conclude that erythroid genes are highly likely to be regulated by enhancers occupied by LDB1 through LDB1 dimerization-mediated looping (Krivega et al., 2014) or by looping mediated by LDB1/CTCF interaction either individually or together (Figure 6B).

## DISCUSSION

The LDB1 transcription complex is the primary mediator of global erythroid gene activation (Fujiwara et al., 2009; Yu et al., 2009; Kassouf et al., 2010; Soler et al., 2010; Li et al., 2013; Mylona et al., 2013). In the best-studied example, the β-globin locus, the complex occupies both the LCR enhancer and β-globin gene. Looping between these elements, upon which transcription activation depends, is mediated by LDB1 dimerization (Deng et al., 2014; Deng et al., 2012; Krivega et al., 2014). Here, we identify numerous LDB1-bound enhancers that interact with erythroid genes occupied by CTCF but not by LDB1 and show, for select genes, that direct interaction between LDB1 and CTCF underlies these contacts. Thus, LDB1 can co-opt CTCF into cell type specific enhancer interactions to contribute to the erythroid transcriptome. This result provides a mechanistic explanation of how LDB1-occupied enhancers can activate genes, such as *Myb* and *Bcl11a*, that are not occupied by LDB1 but do have promoter or first intron CTCF sites.

Recent data have described the differences in the enhancer landscape between neural progenitor cells and erythroid fetal liver cells or the changes observed in the landscape as erythroid progenitor cells differentiate (Schoenfelder et al., 2015; Huang et al., 2016). Here, we focused on the molecular mechanisms employed by enhancers that define the mature erythroid transcriptome. Most (80%) of erythroid genes that loop to at least one enhancer loop to an LDB1-bound enhancer, indicating the predominance of LDB1-mediated enhancer activation of these genes (Li et al., 2013). Of the erythroid genes that loop to these LDB1-bound enhancers and are regulated by LDB1, as defined by reduced expression upon LDB1 KD and rescue upon LDB1 re-expression (Krivega et al., 2014), the largest fraction (39%) were occupied by both LDB1 and CTCF at their promoters. The LDB1 and CTCF binding motifs were almost always (73%) separated by more than 100 bp, suggesting the two kinds of loops are distinct in their gene anchors but may function together (data not shown). We suggest that enhancer/gene loops mediated by more than one mechanism provide stability as well as flexibility to the interactions to influence target gene expression.

An additional 33% of these erythroid, LDB1 sensitive/rescued genes were occupied by CTCF but not LDB1, raising the question whether the looping of these genes to LDB1-occupied enhancers may be independent of LDB1 dimerization. However, transcription of representative genes of this type could not be rescued by expression of LDB1-dimerization deficient mutants (Krivega et al., 2014) in LDB1 KD cells (Figure S6). Interestingly, these genes all looped to multiple LDB1-occupied enhancers. We speculate that LDB1 dimerization may contribute to overall architectural organization required for transcription activation in these loci through looping multiple enhancers together. Enhancer-enhancer looping may be an even stronger component of this organization than enhancer-gene looping (Zhu et al., 2016; Stevens et al., 2017)

We also observed that 198 genes in the ‘other’ set not defined as erythroid are looped to LDB1-bound enhancers and are likely regulated by LDB1, as defined by reduced expression upon LDB1 KD and rescue upon LDB1 re-expression. Although the number is likely to be an overestimate due to the non-exhaustive list of erythroid genes, we suggest that LDB1-bound enhancers have functions beyond those related to erythroid genes. Indeed, recent reports describe LDB1 involvement in long-range gene regulation in select non-erythroid cells (Caputo et al., 2015; Zhang et al., 2015; Costello et al., 2015). Although molecular details remain to be worked out, in these cases LIM only or LIM-homeodomain proteins and DNA binding factors distinct from those in erythroid cells likely mediate LDB1 interaction with DNA. For example, in cardiac progenitor cells, LDB1 can function together with ISL1 to regulate cardiac-specific genes over long distances (Caputo et al., 2015). The potential ability of LDB1 to interact with the large family of tissue specific LIM only and LIM homeodomain proteins, and through them with diverse transcription factors and co-factors, suggests LDB1 is a highly versatile enhancer-looping factor.

Enhancers occupied by LDB1 complexes often coincide with SNP disease-associated genetic variants, according to genome-wide association studies (GWAS) (Maurano et al., 2012; Su et al., 2013). Among these associations are numerous instances where an LDB1 enhancer containing a SNP interacts with a gene relevant to the SNP phenotype whose promoter is occupied by CTCF but not LDB1 (Maurano et al., 2012). In particular, within *Myb* and *Bcl11a* enhancers, SNPs cluster at Ldb1 complex binding sites (Bauer et al., 2013; Stadhouders et al., 2014). Overall, SNPs are known to cluster at looped gene regulatory sites on a genome wide scale (Rao et al., 2014). Thus, understanding the protein players in long-range enhancer looping is likely to suggest targets relevant to multiple genetic diseases.

## EXPERIMENTAL PROCEEDURES

### Cell culture

Mouse erythroid leukemia (MEL) cells and human embryonic kidney (293T) cells were cultured in DMEM and human K562 erythroleukemia cells were cultured in RPMI 1640, with 10% fetal bovine serum in a humidified incubator at 5% CO2. MEL cell differentiation was induced at a concentration of 2.5 × 10^5^ cells per ml with 1.5% DMSO for 4 days.

### Inducible Ldb1 gene deletion in primary cells

Inducible deletion of *Ldb1* in mouse primary erythroid cells from E14.5 embryos was performed as described (Krivega et al., 2014).

### Reporter assay

The Dual Luciferase Reporter Assay system (Promega) and an AutoLumat LB LB 953 luminometer (Berthold) were employed. A 1.57 kb of *Car2* -8/-9 DNA fragment or β-globin LCR HS2 were fused to the SV40 promoter or mouse *Car2* promoter and cloned into the pGL4.2 firefly Luciferase vector. Renilla Luciferase (pGL4.74) served as internal control. Transfections into K562 and MEL cells was performed using Lipofectamine LTX Plus reagent as suggested by the manufacturer (Invitrogen).

### Co-immunoprecipitation (Co-IP)

Co-IP of endogenous proteins was performed using nuclear extracts of MEL cells treated with or without DMSO for 4 days as described (Song et al., 2007). For ectopically expressed proteins, 293T cells were transfected with CTCF-V5 or HA-LDB1. Cells were lysed in IP buffer 1 (25 mM HEPES pH8.0, 500 mM NaCl, 2 mM EDTA, 1 mM NaF, 1 mM NaVO4, 0.1% Tween 20 and protease inhibitors). The cleared lysate was diluted with IP buffer 2 (25 mM HEPES pH8.0, 2 mM EDTA, 1 mM NaF, 1 mM NaVO4, 0.1% Tween 20 and protease inhibitors) to final concentration of 150 mM NaCl. Cells expressing LDB1 deletion mutants were pre-treated with MG132 for 6 hrs to inhibit proteasomal degredation. Cleared extracts were incubated with anti-HA agarose (Sigma) for 2hr at 4°C. The beads were washed with buffer containing 25 mM HEPES pH8.0, 200 mM NaCl, 2 mM EDTA, 1 mM NaF, 1 mM NaVO4, 0.1% Tween 20 and protease inhibitors. Bound proteins were eluted with glycine. Proteins were separated by SDS-PAGE, and Immunoblots were developed with the ECL Plus detection system (Fisher Scientific). For antibodies see Table S2.

### Pull down assay

GST-fused CTCF and MBP-LDB1 proteins were expressed in E. coli BL21(DE3) (Invitrogen) and purified using Glutathione-Sepharose beads (GE Healthcare Inc.) and Amylose resin (New England Biolabs) according to the manufacturer's instructions. Eluted proteins were analyzed by western blot using anti-MBP antibodies (see Table S1).

### Virus Production and Transduction

HA-tagged proteins were constructed with pLenti6/V5-D-TOPO vector (Invitrogen). LDB1 (Clones TRCN0000039019 and -31920) and CTCF (clones TRCN0000096339 and -96340) lentiviral shRNAs were purchased from Open Biosystems. 293FT cells were transduced with vectors and Virapower packaging mix (Invitrogen) according to the manufacturer’s instructions except that high speed centrifugation was performed on a cushion of 20% Sucrose. For stable clones and pools, MEL cells were incubated with viral particles in the presence of 6 ug/ml polybrene and selected with 5μg of Puromycin or 40 μg of Blasticidin for up to 2 weeks.

### Chromatin immunoprecipitation (ChIP)

ChIP was performed as described (Song et al., 2007). Differences in DNA enrichment were determined by real-time qPCR using SYBR chemistry with the ABI 7900HT (Applied Biosystems). Results from 3 independent replicates are shown and error bars indicate SEM. For antibodies see Table S2. For ChIP primers see Table S3.

### Chromatin conformation capture assay (3C)

3C was performed as described (Tolhuis et al., 2002; Song et al., 2007; Hagege et al., 2007), except that formaldehyde-crosslinked chromatin was digested twice with BstYI or EcoRI overnight. Digestion efficiency was monitored as described (Hagege et al., 2007). Bacterial artificial chromosomes (BACs) containing the CAR (RP23-330N22 and RP24-385I21) or Cpeb4 (BMQ-72D14) loci and ERCC3 (RP24-97P16) were digested with BstYI or EcoRI and religated to monitor PCR efficiency (control template). Real time qPCR was carried out on an ABI 7900HT instrument using Taqman probes and primers (Table S4). Values were normalized to ERCC3 (Palstra et al., 2003) and to 2 non-interacting fragments outside of *Car2/Car3*. Results from 3 independent replicates are shown and error bars indicate SEM.

### Reverse-Transcription Reaction

DNase I treated RNA (1 ug) was reverse transcribed by using the Superscript III according to the manufacturer (Invitrogen). cDNA was diluted to 200 ul, and 2 ul of cDNA was amplified in a 10 ul or 25 μl reaction volume by real-time qPCR by using SYBR chemistry. For primers see Table S3. Results from at least 3 different RNA preparations are shown and data were normalized to Gapdh. Error bars in the figures represent SEM.

### Western blotting

Cells were lysed in RIPA buffer (50 mM Tris-HCl at pH 8, 150 mM NaCl, 1% NP-40, 0.5% Na deoxycholate, 0.1% SDS) and protein concentration determined using BCA protein assay kit (Thermo Scientific). Sample were separated by NuPAGE gel, transfered to PVDF membranes according to the manufacturer’s instructions, and probed with antibodies listed in Table S1. Blots were developed by ECL Plus (Thermo Scientific).

### Gene editing using CRISPR/Cas9

MEL cells were transfected with Lipofectamine LTX plus (Invitrogen) or Nucleofactor (Lonza) following manufacturers suggestions. Cas9 expression vector (pCas9_GFP, #44720, a gift of K. Musunuru) and guide RNA expression vectors (pgDNA) were obtained from Addgene. Targeted sequences are shown in Table S4. 48hr post-transfection, the top 0.1 % of EGFP positive cells were sorted by FACS ARIA II (BD biosciences) and individual clones were grown in 96 well plates. Genomic DNA was purified with DNeasy blood and tissue kit (Qiagen) and genotyping was performed with Q5 taq polymerase (NEB) and target specific primers flanking the deletions (Figure S1). Deletions were validated by sequencing (Table S1).

### Computational methods, data acquisition and preparation

See Supplementary Materials and Methods for details.

## ACKNOWLEDGEMENTS

We thank Dr. Gary Felsenfeld for the GST-CTCF expression vectors. We acknowledge the NIDDK Sequencing Core for sequencing. This work was funded by the Intramural Program of the National Institute of Diabetes, Digestive and Kidney Diseases, NIH (DK015508 to A.D.).

## AUTHOR CONTRIBUTIONS

J.L. and A.D. conceived the project. J.L. and I.K. conducted experiments. J.L., R.K.D., I.K. and A.D. analyzed data and wrote the paper. The authors declare no conflicts of interest.

